# Intracellular BH3 profiling reveals shifts in anti-apoptotic dependency in B-cell maturation and activation

**DOI:** 10.1101/148437

**Authors:** Joanne Dai, Micah A. Luftig

## Abstract

Apoptosis is critical to B-cell maturation, but studies of apoptotic regulation in primary human B cells is lacking. Previously, we found that infecting human B cells with Epstein-Barr virus induces two different survival strategies (Price *et al.*, 2017). Here, we sought to better understand the mechanisms of apoptotic regulation in normal and activated B cells. Using intracellular BH3 profiling (iBH3), we defined the Bcl2-dependency of B-cell subsets from human peripheral blood and tonsillar lymphoid tissue as well as mitogen-activated B cells. We found that naïve and memory B cells were BCL-2 dependent, while germinal center B cells were MCL-1 dependent and plasma cells were BCL-XL dependent. Proliferating B cells activated by CpG or CD40L/IL-4 became more dependent upon MCL-1 and BCL-XL. As B-cell lymphomas often rely on survival mechanisms derived from normal and activated B cells, these findings offer new insight into potential therapeutic strategies for lymphomas.

## Introduction

The maturation of B cells into antibody-secreting cells is critical to immunity and is accomplished through either T-independent or T-dependent B-cell maturation. In T-independent B-cell maturation, antigen binding to the B-cell receptor (BCR) stimulates extra-follicular naïve and memory B cells to proliferate (Bortnick and Allman, 2013). B cells can also be activated via pathogen-associated molecular patterns (PAMPs), like CpG DNA, binding to Toll-like receptors (TLR) (Bernasconi et al., 2002; Krieg et al., 1995). Activated B cells can differentiate into short-lived plasmablasts or migrate into secondary lymphoid tissue to undergo T-dependent maturation. In the germinal center (GC) reaction, the affinity for antigen is further refined through somatic hypermutation and selection by apoptosis (Victora and Nussenzweig, 2012). B cells also compete for survival signals in the form of antigen binding to the BCR and CD40 receptor binding by to cognate T follicular helper cells. Outcompeted B cells are eliminated by apoptosis and surviving B cells exit the GC as long-lived memory B cells or plasma cells.

Ultimately, the humoral repertoire is shaped by apoptosis dependent on the multi-domain Bcl-2 family proteins that select which B cells die and which continue to propagate (Peperzak et al., 2012). Apoptosis is initiated when BAX and BAK dimerize and permeabilize the mitochondrial outer membrane to release cytochrome C into the cytosol, which activates a cascade of caspases that mediate DNA fragmentation, protein degradation, and cellular blebbing (Fischer et al., 2003). To prevent aberrant activation, anti-apoptotic proteins bind and sequester BAX and BAK as well as the pro-apoptotic BH3-only proteins (named as such because they share homology in their BH3 domain). Dysregulation of apoptosis typically accompanies the development of lymphomas or autoimmunity, and understanding the mechanisms of apoptosis regulation is key to provide insight and develop new therapeutic strategies for these diseases (Vo et al., 2012).

New breakthroughs in studying apoptosis regulation during B-cell maturation have been made possible by the development of BH3 mimetics, a class of small molecular inhibitors that mimic pro-apoptotic proteins and have shown great promise as cancer therapeutics (Oltersdorf et al., 2005). In mice, naïve and memory B cells are sensitive to the BH3 mimetic ABT-737, which inhibits BCL-2, BCL-XL, and BCL-W, but pre-existing GC and plasma cells are resistant (Carrington et al., 2010), suggesting that apoptosis is regulated differently at various stages of B-cell maturation. Recently, human tonsillar GC B cells were found to be sensitive to MCL-1 inhibition, which supports previous findings that MCL-1 is essential for GC formation in mice (Peperzak et al., 2017; Vikstrom et al., 2010). Nonetheless, there remain challenges associated with studying apoptosis in primary human B cells, which have short life-spans in culture, spontaneously undergo apoptosis, and are difficult to transfect. In addition, how apoptotic regulation changes specifically during B-cell activation poorly understood. As a result, we utilized intracellular BH3 (iBH3) profiling to uncover the dominant anti-apoptotic mechanism at the mitochondria of resting and activated primary human B cells. The principles behind iBH3 profiling are based upon the selective interactions between the pro- and anti-apoptotic proteins of the Bcl2-family (Deng et al., 2007; Youle and Strasser, 2008). In short, the anti-apoptotic proteins that are present at the mitochondria dictate which pro-apoptotic BH3-only peptides can induce mitochondrial depolarization and cytochrome C release. In this way, we can identify the anti-apoptotic dependency, or Bcl2-depedency, of the sample. iBH3 profiling can also assay the Bcl2-dependency of multiple populations in heterogeneous samples, thereby eliminating the need for time-consuming and expensive sorting experiments to study rare populations (Ryan et al., 2016).

With the advent of iBH3 profiling and BH3 mimetics, we are equipped with tools that allow us to study how apoptosis is regulated in human B-cell subsets and activated B cells. Therefore, we have used iBH3 profiling to determine the BH3 profiles of normal, resting, and activated B cells, and validated the hypotheses generated by these profiles by treating normal and activated human B-cell subsets with BH3 mimetics.

## Results

### Naïve and memory B cells circulating in the periphery depend upon BCL-2 for survival

To ensure the reliability of iBH3 profiling, we sought to determine the mechanism of apoptosis regulation in peripheral blood B cells from within the bulk PBMC population (Carrington et al., 2010). After staining for cell-specific surface markers, cells were incubated with recombinant BH3-only peptides and digitonin, which selectively permeabilizes the outer cellular membrane and exposes intact mitochondria to the peptide treatment (**Figure 1A**). Following peptide treatment, the cells were fixed, incubated with saponin to permeabilize mitochondria, and stained with a fluorescently-labeled antibody to quantify the remaining levels of intracellular cytochrome C. The extent of cytochrome C release was measured as the dynamic range between total levels of cytochrome C, determined by treating with an inert recombinant PUMA BH3-only peptide (PUMA2A) and alamethicin (Alm), which can induce 100% cytochrome C release.

**Figure 1.**
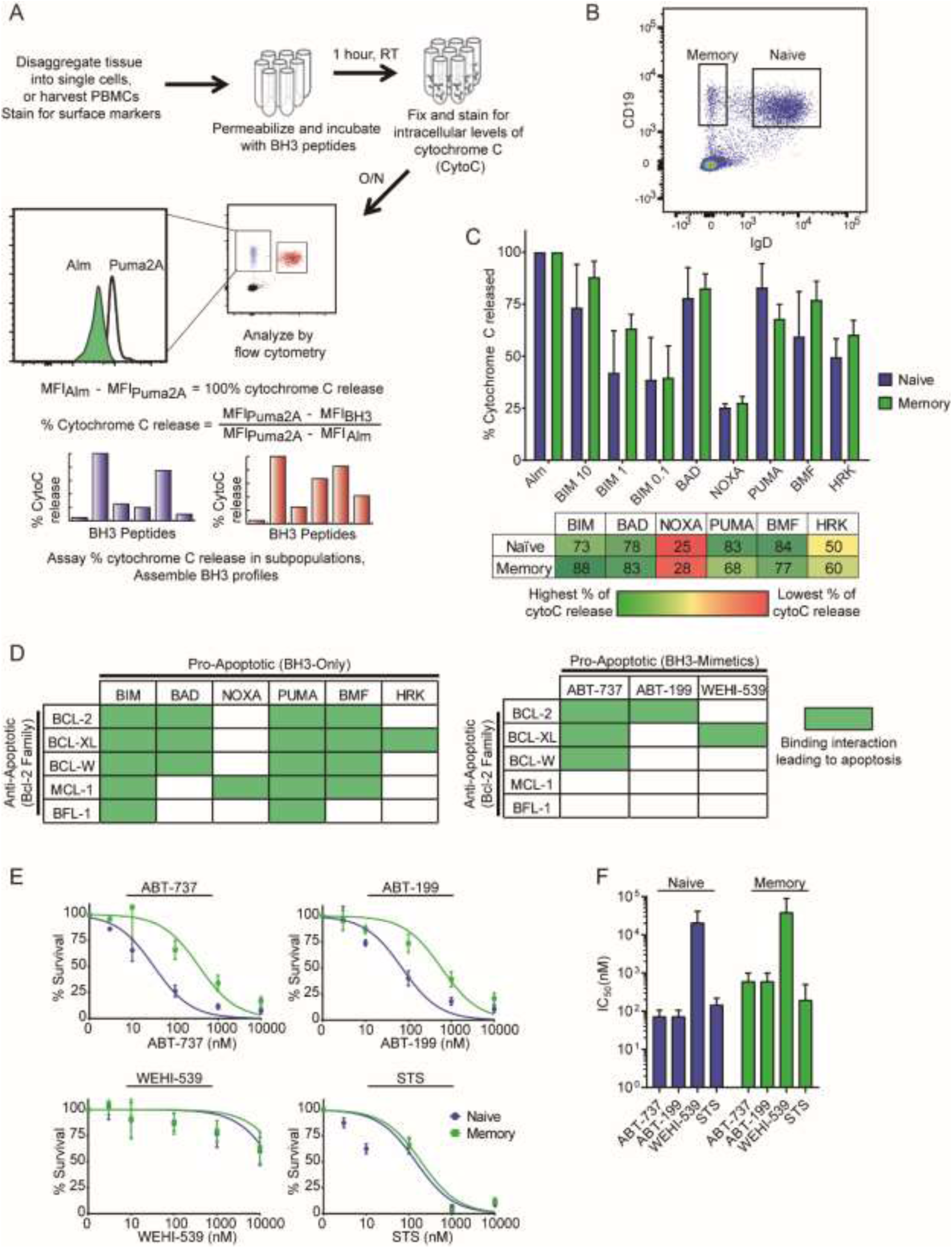
The iBH3 profiles of normal, resting human B cells from the peripheral blood. A) Schematic of the intracellular BH3 (iBH3) process, including how percentage of cytochrome C release is quantified. B) Representative flow cytometry plot of CD19^+^ naïve (IgD^+^) and memory B cells (IgD^-^) from human peripheral blood mononuclear cells (PBMCs). C) iBH3 profiles of naïve and memory B cells from three human donors. Mean and SEM are plotted; mean values are summarized in table below. Percentage of cytochrome C release is normalized to alamethicin (Alm) positive control and Puma2A negative control. BIM 10 - 10μM BIM, BIM 1 - 1μM BIM, BIM 0.1 – 0.1μM BIM D) Left, schematic of selective interactions between pro- and anti-apoptotic members of the Bcl-2 protein family and, right, BH3 mimetics and the antiapoptotic proteins that they inhibit. Green boxes indicate a binding interaction between the two proteins that leads to cytochrome C release. E) Dose-response curves generated from treating PBMCs for 24 hours with ABT-737 (BCL-2, BCL-XL, BCL-W inhibitor; 8 human donors), ABT-199 (BCL-2 inhibitor; 8 human donors), WEHI-539 (BCL-XL inhibitor; 3 human donors) and staurosporine (STS; Naïve, 9 human donors; Memory, 8 human donors), a broad kinase inhibitor that can induce apoptosis. Percent survival is measured as a percentage of DMSO-treated controls. F) Average IC_50_ values with 95% confidence intervals for naïve and memory B cells are plotted from (E).

Naïve (CD19^+^ IgD^+^) and memory (CD19^+^ IgD^-^) B cells were distinguished by IgD expression (**Figure 1B**). iBH3 profiling revealed that the mitochondria in naïve and memory B cells in the peripheral blood were highly sensitive to BIM, BAD, PUMA, and BMF, and relatively resistant to NOXA and HRK (**Figure 1C**). This pattern of sensitivity suggested that naïve and memory B cells were dependent upon BCL-2 for survival (**Figure 1D**).

To validate this hypothesis, we treated PBMCs with the following BH3 mimetics: ABT-737 (BCL-2, BCL-XL, BCL-W inhibitor), ABT-199 (BCL-2 inhibitor), or WEHI-539 (BCL-XL inhibitor) (**Figure 1E**). Naïve and memory B cells were resistant to WEHI-539 (IC_50_ >1 μM) and were sensitive to ABT-737 and ABT-199 (IC_50_ <1 μM), thereby confirming our iBH3 profiling hypotheses and previously published results of BCL-2 dependency (**Figure 1F**). ABT-199 was used to distinguish between BCL-2 and BCL-W, which have identical binding partners (**Figure 1D**). Staurosporine (STS) is not a BH3 mimetic, but rather it is a pan-kinase inhibitor that induces apoptosis and was used here to distinguish cells that can undergo apoptosis from those that can’t, such as cells that lack the effector molecules BAX and BAK (Wei et al., 2001).

### B-cell maturation is associated with changes in Bcl-2 family dependency

To determine the changes in apoptotic regulation that take place during human B-cell maturation, we queried the apoptotic regulation of CD19^+^ naïve (IgD^+^ CD38^-^), memory (IgD^-^ CD38^-^), germinal center (GC; IgD^-^ CD38^mid^) B cells, and plasma cells (IgD^-^ CD38^hi^) from tonsillar tissue (**Figure 2A**). Like naïve and memory B cells in peripheral blood, tonsillar naïve and memory B cells were highly sensitive to BIM, BAD, PUMA, and BMF and were predicted to be BCL-2 dependent (**Figure 2B-C**). GC B cells, however, had a modestly lower response to BAD and a heightened response to NOXA, suggesting that GC B cells were dependent upon MCL-1 for survival. Plasma cells were much less responsive to BH3 peptide treatments compared to the other B-cell subsets. However, we hypothesized that within the reduced dynamic range of plasma cell responses, lower NOXA sensitivity as compared to BAD and HRK suggested dependence on BCL-XL for survival.

**Figure 2.**
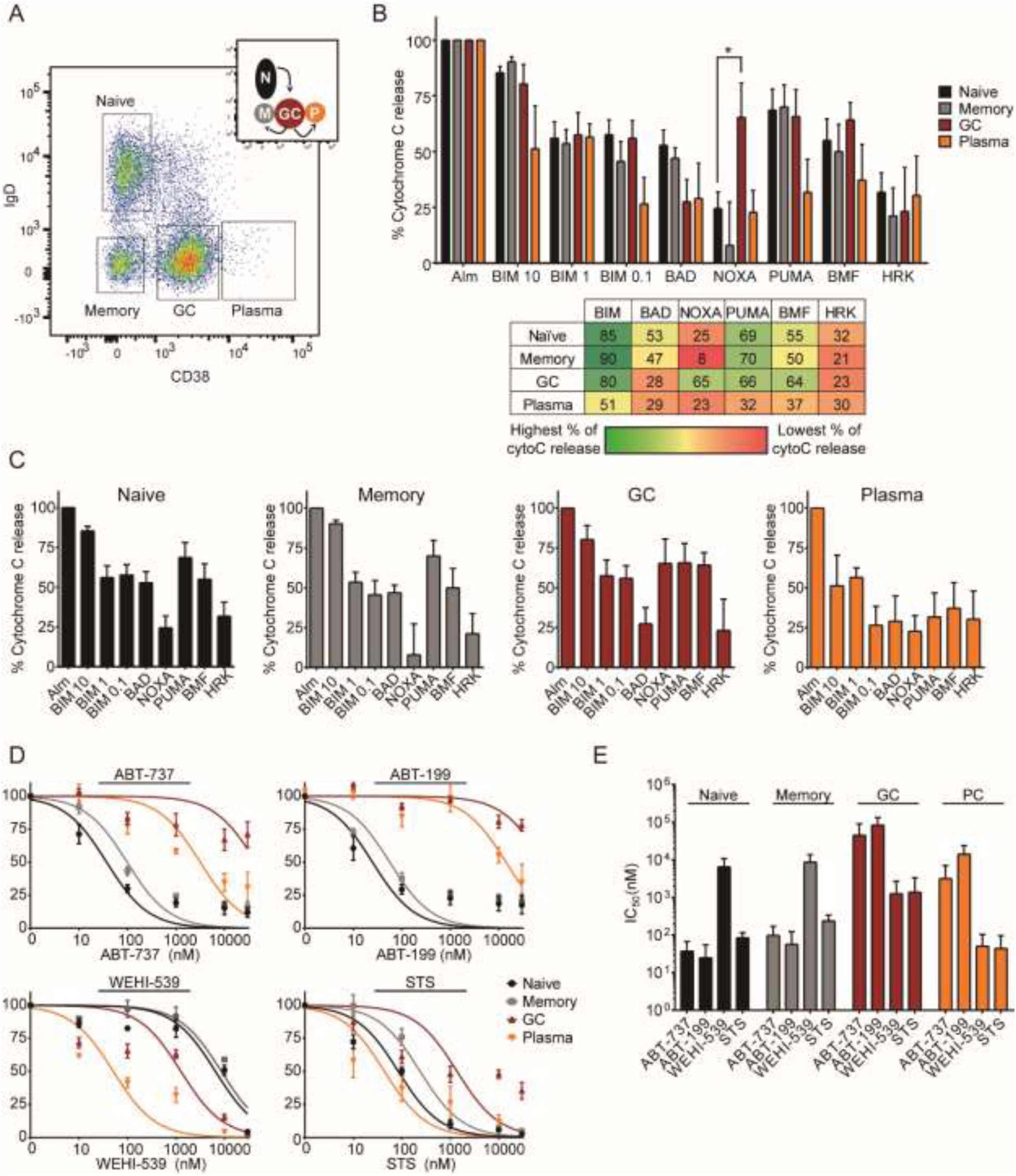
The iBH3 profiles of normal, resting human B cells from tonsillar lymphoid tissue. A) Representative flow cytometry plot of CD19^+^ B cell subsets found in human tonsillar tissue: naïve (IgD^+^ CD38^-^), memory (IgD^-^ CD38^-^), germinal center (GC; IgD^-^ CD38^mid^), and plasma cells (IgD^-^ CD38^hi^). Inset, a color-coded schematic of the GC reaction. B) iBH3 profiles of tonsillar B cell subsets from 4-5 human donors. Mean and SEM are plotted; mean values are summarized in the table below. Percentage of cytochrome C release is normalized to alamethicin (Alm) positive control and Puma2A negative control. In samples treated with Noxa-derived recombinant peptides, * p=0.0467, by unpaired two-tailed Student’s t-test. C) Same as in (B), but showing each subset’s profile individually. D) Dose-response curves generated from treating eight human donors for 24 hours with ABT-737 (BCL-2, BCL-XL, BCL-W inhibitor; 3 human donors), ABT-199 (BCL-2 inhibitor; 3 human donors), WEHI-539 (BCL-XL inhibitor; 3 human donors) and staurosporine (STS; 4-6 human donors), a broad kinase inhibitor that can induce apoptosis. Percent survival is measured as a percentage of DMSO-treated controls. E) Average IC_50_ values with 95% confidence intervals for each subset are plotted from (D). Plasma cells, PC.

To validate the hypotheses generated from iBH3 profiling, we treated primary tonsillar B cells with ABT-199, ABT-737, WEHI-539, and STS (**Figure 2D-E**). All cells were sensitive to STS, indicating that all subsets were capable of intrinsic apoptosis. As predicted, naïve and memory B cells were sensitive to BCL-2 inhibition by ABT-199 and ABT-737. Plasma cells were uniquely sensitive to BCL-XL inhibition by WEHI-539, confirming their dependence on BCL-XL. The difference in the sensitivity of plasma cells to ABT-737 and WEHI-539 may be due to a difference in their binding affinities for BCL-XL (ABT-737 IC_50_=35nM vs WEHI-539 IC_50_=1.1nM) (Lessene et al., 2013; Oltersdorf et al., 2005). GC B cells were resistant to ABT-737, ABT-199, and WEHI-539 since none of these drugs inhibited MCL-1. However, GC B cells were also resistant to the MCL-1 specific BH3 mimetic, A-1210477 (**Figure 2 Figure Supplement 1**), which is likely due to its weak cellular potency (i.e. sensitivity in the micromolar range) and nonspecific toxicity (Kotschy et al., 2016; Leverson et al., 2015). In sum, these findings support an existing model that B cells adjust their Bcl-2 family dependency throughout T-dependent B cell maturation. Moreover, iBH3 profiling proved to be reliable in querying heterogeneous populations simultaneously.

### CD40L/IL-4 stimulated B cells are sensitive to NOXA and HRK peptides, yet remain sensitive to BCL-2 inhibition

We next sought to model B-cell maturation with mitogen stimulation of peripheral blood B cells. To track proliferation, PBMCs were stained with CellTrace Violet (CTV), which becomes diluted upon cell division (**Figure 3A**). CTV was retained during permeabilization with digitonin, thereby making it suitable in iBH3 profiling. We induced proliferation in peripheral blood B cells *ex vivo* by treating with CD40L and IL-4 to mimic T-cell derived signals important in B-cell survival in the GC. CD19+ B cells began proliferating 3 days after treatment with CD40L/IL-4, as shown previously (Hawkins et al., 2007; Nikitin et al., 2014).

**Figure 3.**
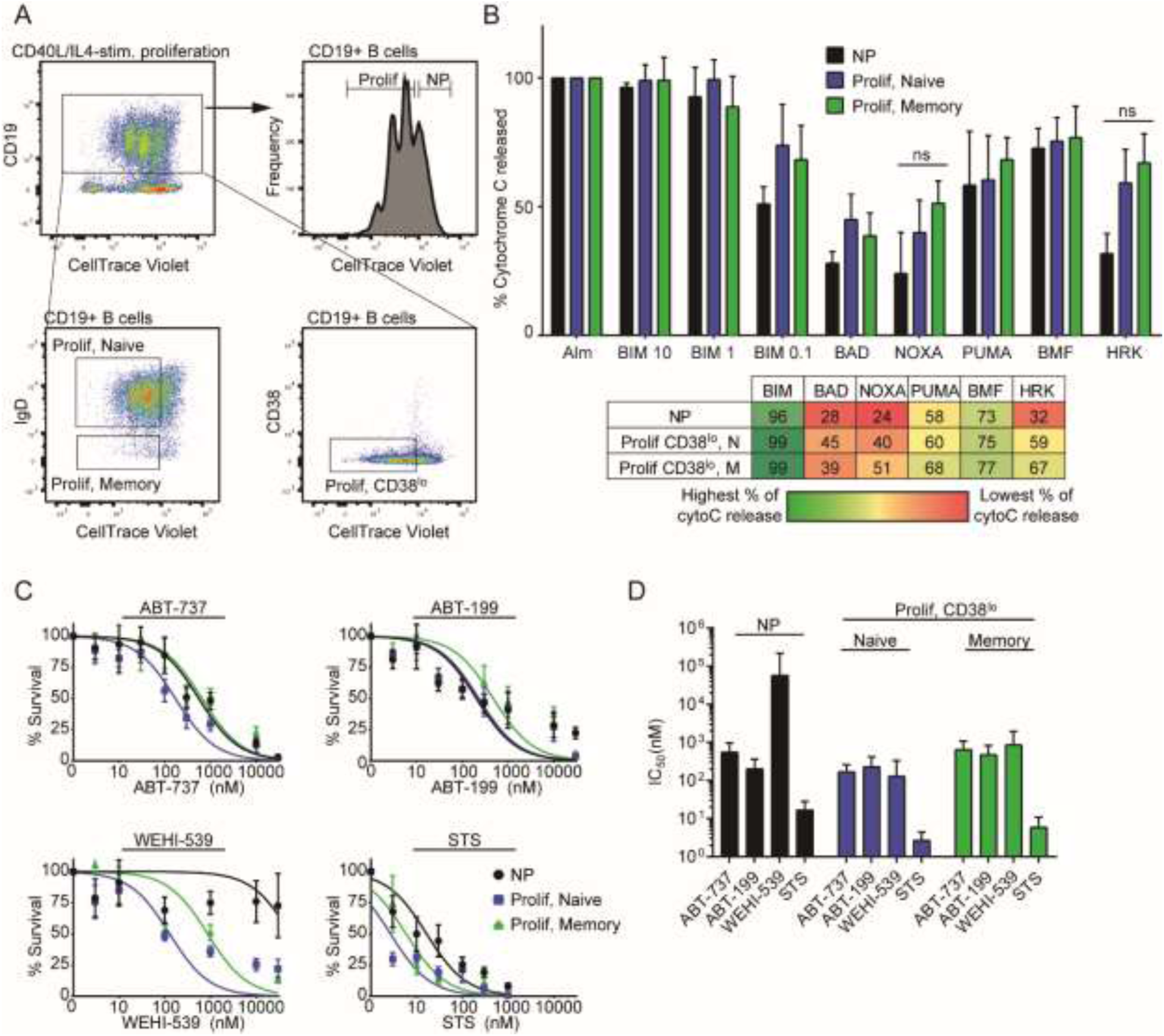
Changes in apoptotic regulation in activated, proliferating B cells stimulated by CD40L/IL-4. A) Representative flow cytometry plot of CD19^+^ B cells that have been stimulated to proliferate by CD40L/IL-4, Day 7 after stimulation. Dilution of CellTrace Violet with each division distinguishes proliferating (Prolif; CTV^lo^) cells from nonproliferating (NP; CTV^hi^) cells. Cells that have proliferated the furthest also upregulate CD38 surface expression to some extent, but the size of this population has been inconsistent across donors. Both naïve and memory B cells proliferate in response to CD40L/IL-4 stimulation. B) iBH3 profiles of different subsets of CD40L/IL-4 stimulated B cells from 3-5 human donors. Mean and SEM are plotted; mean values are summarized in the table below. Percentage of cytochrome C release is normalized to alamethicin positive control and Puma2A negative control. ns, not significant by unpaired two-tailed Student’s t-test. C) Dose-response curves generated from treating three human donors for 3 days with ABT-737 (6 human donors), ABT-199 (4-9 human donors), WEHI-539 (3 human donors), and STS (6 human donors). D) Average IC_50_ values with 95% confidence intervals for each subset are plotted from (C).

CD40L/IL-4 stimulated B cells were iBH3 profiled six days post-stimulation (**Figure 3B**). Proliferating B cells displayed modestly increased sensitivity to NOXA and HRK peptide treatment relative to nonproliferating B cells, suggesting an increasing dependence on MCL-1 and BCL-XL for survival. The robust increase in HRK sensitivity correlated with increased susceptibility of proliferating B cells to the BCL-XL inhibitor, WEHI-539 (**Figure 3C-D**). However, these cells remained sensitive to ABT-199 and ABT-737, suggesting that BCL-2 still plays an important role in CD40L/IL-4 stimulated B-cell survival (**Figure 3C-D**). This also indicates that NOXA sensitivity observed in proliferating B cells did not confer MCL-1 dependence.

### A CD38^hi^, plasma blast-like subpopulation in CpG-stimulated B cells becomes sensitive to NOXA and HRK in iBH3 profiling and becomes resistant to BCL-2 inhibition

In B cells activated to proliferate *ex vivo* by CpG DNA, a PAMP that stimulates the TLR9 pathway, we observed similar changes in apoptotic regulation in proliferating B cells. However, in CpG-stimulated cells, B cells that have undergone the most cellular divisions upregulate high levels of CD38 surface expression, suggesting that they have differentiated into plasmablast-like B cells (CTV^lo^ CD38^hi^; **Figure 4A**) (Bernasconi et al., 2002; Huggins et al., 2007). This population consisted predominantly of memory B cells and is noticeably absent in CD40L/IL-4 stimulated B cells because IL-4 downregulates CD38 expression through serine/threonine kinases (Shubinsky and Schlesinger, 1996). In iBH3 profiling of CpG-stimulated B cells, nonproliferating B cells were predicted to be BCL-2 dependent and as they proliferate they become more responsive to NOXA and HRK peptides with CD38 ^hi^ B cells as the most responsive to NOXA and HRK (**Figure 4B**).

**Figure 4.**
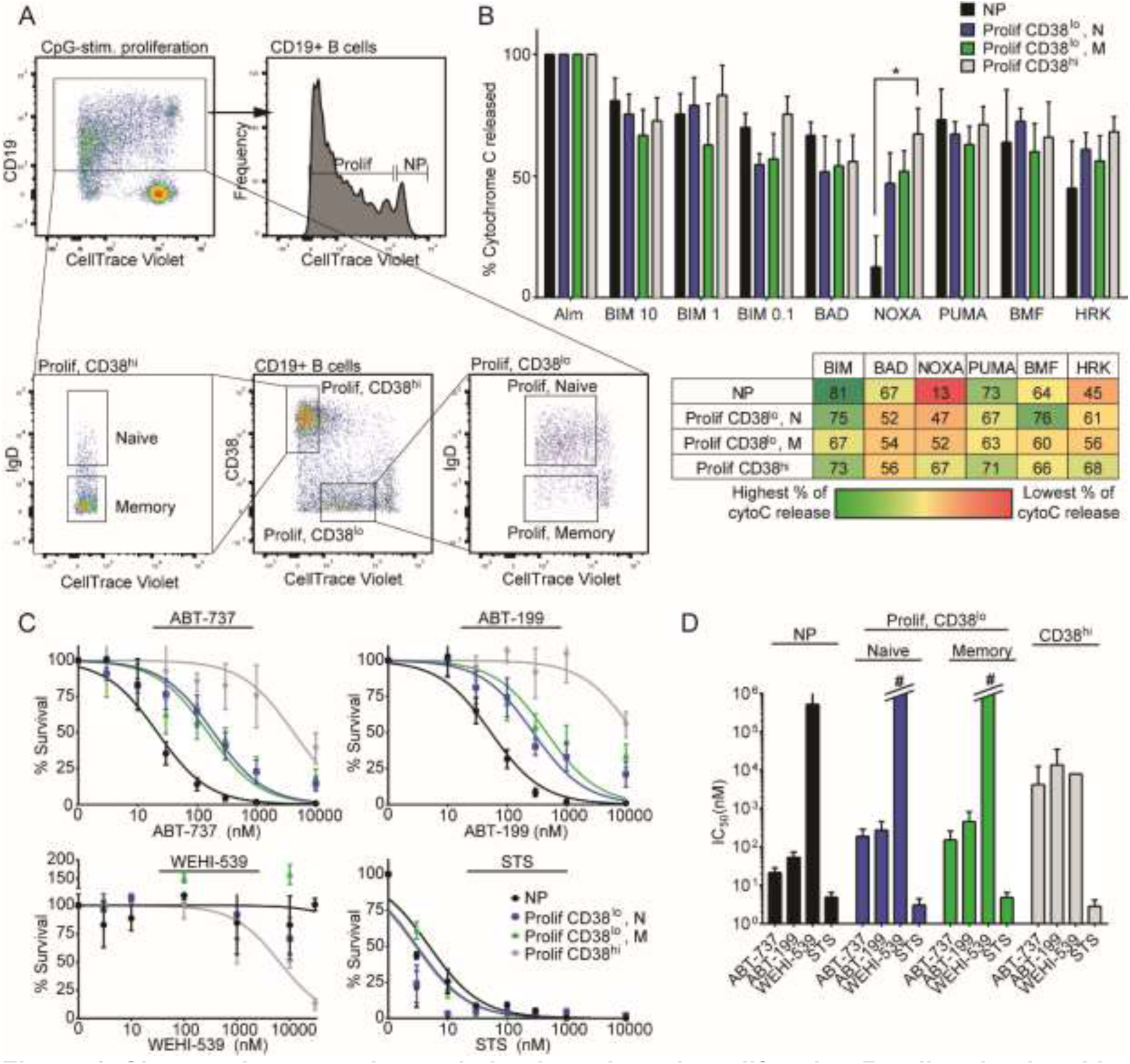
Changes in apoptotic regulation in activated, proliferating B cells stimulated by CpG. A) Representative flow cytometry plot of CD19^+^ B cells that have been stimulated to proliferate by CpG, Day 6 after stimulation. Dilution of CellTrace Violet with each division distinguishes proliferating (Prolif) cells from nonproliferating (NP) cells. Cells that have proliferated the most also upregulate high CD38 surface expression (CD38^hi^). Both naïve and memory B cells proliferate in response to CpG stimulation. B) iBH3 profiles of different subsets of CpG-stimulated B cells from 4 human donors. Mean and SEM are plotted; mean values are summarized in the table below. Percentage of cytochrome C release is normalized to alamethicin (Alm) positive control and Puma2A negative control. In samples treated with Noxaderived peptide, * p=0.0292, by unpaired two-tailed Student’s t-test. C) Dose-response curves generated from CpG-stimulated PBMCs for 3 days with ABT-737 (5-7 human donors), ABT-199 (5-7 human donors), WEHI-539 (3 human donors), and STS (6-7 human donors). D) Average IC_50_ values with 95% confidence intervals for each subset are plotted from (C). #, for these subsets, there is no detectable sensitivity and therefore no calculable IC_50_ value.

Nonproliferating and CD38^lo^ proliferating B cells were within the same range of sensitivity to ABT-199 and ABT-737 and were refractory to WEHI-539, indicating that as B cells proliferated in response to CpG, BCL-2 remained important in regulating survival (**Figure 4C-D**). CD38^hi^ B cells, in contrast, were less sensitive to ABT-199, ABT-737, and slightly more sensitive to WEHI-539. Interestingly, it appeared that WEHI-539 increased the number of proliferating CD38^lo^ memory B cells (**Figure 4C**), suggesting that in T-independent B-cell maturation, BCLXL may be involved in the differentiation of plasmablasts derived from memory B cells. This also indicates that the perceived sensitivity of the CD38^hi^ cells to WEHI-539 may not be due to cell death, but rather from a block in differentiation.

The overall increase in apoptosis resistance in the CD38^hi^ subset is not because of impaired BAX and BAK activation, since CD38^hi^ cells were sensitive to STS, but rather it may be due to an increasing dependence upon MCL-1 as indicated by increasing sensitivity to NOXA in their iBH3 profile. In fact, of the proliferating subsets, only the response of the CD38^hi^ subset to NOXA was significantly different from that of nonproliferating B cells (* p=0.0292). Thus, to inhibit both BCL-XL and MCL-1, we tested the efficacy of combining WEHI-539 with A-1210477 in CpG-stimulated PBMCs (Chou, 2010). CpG-stimulated B cells were refractory to high concentrations of A-1210477 (**Figure 4 Figure Supplement 1A-B**). However, a combination of A-1210477 and sub-IC50 levels of WEHI-539 resulted in synergism (Combination Index, CI<1) in the CD38^hi^ subpopulation (**Figure 4 Figure Supplement 1C**). This effect was specific to MCL-1 as ABT-199 did not synergize with WEHI-539 (CI≈1) (**Figure 4 Figure Supplement 1D**).

## Discussion

Proper regulation of apoptosis is critical for B-lymphocyte development and maturation. Much of our mechanistic understanding of B-cell apoptosis is derived from murine studies due to technical limitations in studying B-cell subsets in humans (Carrington et al., 2010; Peperzak et al., 2013; Vikstrom et al., 2010). While two recent studies have characterized Bcl-2 family dependency in human primary and secondary lymphoid tissue (Peperzak et al., 2017; Sarosiek et al., 2017), in this study we use the newly developed iBH3 profiling technique to query the mechanisms underlying apoptotic regulation both in heterogeneous lymphoid tissue as well as upon activation of normal B-cell subsets *ex vivo* (Ryan and Letai, 2013; Ryan et al., 2016). Our experimental approach couples proliferation tracking with cell surface markers to measurement of intracellular cytochrome C release in response to BH3 peptides to define Bcl2 family member dependency at the single cell level during B-cell activation. Validation of iBH3 profiling with BH3 mimetics defined transitions during B-cell maturation from BCL-2 dependency in naïve and memory B cells, to MCL-1 in GC B cells, and to BCL-XL in plasma cells from secondary lymphoid tissue. Then, upon activation by CpG or CD40L/IL-4, the Bcl2-dependency of proliferating B cells shifts from BCL-2 to a combination of BCL-XL and MCL-1. This shift is apparent by the increased resistance to BCL2-specific BH3 mimetics and is most pronounced in the most differentiated subset of CpG-stimulated B cells.

Our findings corroborate several *in vivo* studies in mice that suggest that stage-specific Bcl2-dependency occurs throughout B-cell maturation. Our work indicates that human naïve and memory B cells, like those in mice, are sensitive to BCL-2 inhibition, whereas GC and plasma cells are refractory (Carrington et al., 2010). In mouse GC B cells or those induced to differentiate into plasma cells *ex vivo*, MCL-1 is important in promoting survival (Peperzak et al., 2013; Vikstrom et al., 2010). Furthermore, murine plasma cells depend on BCL-XL protection from apoptosis due to the unfolded protein response (Gaudette et al., 2014). These parallels in mouse studies with our findings in human B cells suggest that intrinsic apoptotic regulation in B-cell maturation is highly conserved and functionally critical to humoral immunity.

Proliferating B cells in the GC as well as *ex vivo* mitogen-stimulated B blasts display evidence of oxidative and metabolic stress as well as DNA damage, which if left unresolved can lead to apoptosis (Jellusova et al., 2017; Nikitin et al., 2014; Ranuncolo et al., 2007; Wheeler and Defranco, 2012). Our findings indicate that GC cells display a heightened response to NOXA in iBH3 profiling suggesting an increased dependence on MCL-1. In addition, our lab has found that following EBV infection of primary human B cells, which is characterized by hyperproliferation and activation of the DNA damage response (Nikitin et al., 2010), there is a shift in Bcl2-dependency from BCL-2 to MCL-1 (Price et al., 2017). This suggests that upon proliferation, MCL-1 expression is rapidly induced to respond to apoptosis-inducing intracellular stress resulting from rapid expansion and differentiation.

Constitutive activation of gene expression and survival programs found in normal B-cell maturation can promote the development of lymphomas. For example, in follicular lymphoma, a t(14;18) chromosomal translocation induces BCL-2 over-expression in naïve B cells thereby promoting survival in the B-cell follicle of a GC (Kridel et al., 2012; Tsujimoto et al., 1985). Consistent with our findings and that of others linking GC survival to MCL-1 up-regulation (Peperzak et al., 2017; Vikstrom et al., 2010), a subset of DLBCLs that arise from the GC often display MCL-1 copy number gains (Wenzel et al., 2013). Indeed, c-Myc-driven Burkitt’s lymphomas are thought to have arisen from GC B cells, express high levels of MCL-1 and are sensitive to MCL-1 inhibition (Dave et al., 2006; Kelly et al., 2014; Klein et al., 1995; Kotschy et al., 2016). MCL-1 is also often over-expressed in plasma-cell derived multiple myelomas, mimicking the requisite up-regulation of MCL-1 induced by stromal interactions in the bone marrow (Gupta et al., 2017; Wuilleme-Toumi et al., 2005). Interestingly, a subset of plasma-cell derived tumors also display co-dependence of MCL-1 and BCL-XL (Morales et al., 2011), similar to our findings in CpG-induced CD38^hi^ plasmablasts. Overall, these data strongly support the importance of defining normal and activated human B-cell survival mechanisms towards understanding the underlying basis of B-lymphoma cell survival.

## Materials and Methods

### Cells and mitogens

Peripheral B cells were obtained from normal human donors through the Gulf Coast Regional Blood Center (Houston, TX). Buffy coats were layered over Ficoll Histopaque-1077 gradient (Sigma, H8889) and washed three times with FACS buffer (5% FBS in PBS) and cultured in RPMI supplemented with 15% FBS (Corning), 2 mM L-Glutamine, 100 U/ml penicillin, 100 μg/ml streptomycin (Invitrogen), and cyclosporin A. To track proliferation, PBMCs were stained with CellTrace Violet (Invitrogen, C34557) for 20min in PBS at 37°C, washed in FACS buffer, and then cultured with the appropriate mitogens.

Tonsillar B cell subsets were obtained from discarded, anonymized tonsillectomies from the Duke Biospecimen Repository and Processing Core (Durham, NC). Tonsillectomies were manually disaggregated, filtered through a cell strainer, and isolated by Ficoll Histopaque-1077 (Sigma, H8889). The harvested lymphocyte layer was washed three times with FACS buffer and cultured in RPMI supplemented with 15% FBS (Corning), 2 mM L-Glutamine, 100 U/ml penicillin, 100 μg/ml streptomycin (Invitrogen), and cyclosporin A.

TLR9 ligand CpG oligonucleotide (ODN 2006) was purchased from IDT and used at 2.5 μg/mL (Krieg et al., 1995). Human recombinant interleukin-4 (IL-4; Peprotech, AF200-04) was used at 20ng/mL. HA-tagged CD40 ligand (R&D Systems, 6420-CL) was used at 5ng/mL with an anti-HA cross-linking peptide (R&D Systems, MAB060; RRID:AB_10719128) at a concentration of 0.2 μg/μL.

### Intracellular BH3 profiling and validation

Intracellular BH3 (iBH3) profiling was performed with recombinant peptides synthesized by New England Peptides (sequences listed in table). Stock peptides were resuspended at 10mM in DMSO and aliquoted and stored at −80°C. To prepare peptides for iBH3 profiling, peptides were diluted to 200μM in 0.004% digitonin in DTEB buffer (135mM trehalose (Sigma, T9449), 20μM EDTA (Sigma, E6758), 20μM EGTA (Sigma, E3889), 5mM succinic acid (Sigma, S3674), 0.1% IgG-free BSA (VWR, 100182-742), 10mM HEPES (Sigma H4034), 50mM potassium chloride (Sigma, P9541), adjusted to pH 7.5 with potassium hydroxide). Bim peptides were used at 10μM, 1μM, and 0.1μM Alamethicin (Enzo, BML-A150-0005) was used at 25μM.

PBMCs and tonsillar cells were iBH3 profiled immediately after processing, while mitogen-stimulated cells were profiled on day 6 post-stimulation. To prepare samples for iBH3 profiling, cells were stained with the Zombie Aqua viability dye (Biolegend, 423101) in serum-free PBS for 15min at room temperature and then stained with FACS antibodies for surface markers for 30min at 4°C. Because Zombie Aqua and CellTrace Violet fluoresce in the same channel, mitogen-stimulated cells were centrifuged at low speed and washed to remove dead cells (ie Trypan positive) before staining. After staining, cells were pelleted and resuspended at 4 × 10^6^ cells/mL in DTEB buffer. Equal volumes (50μL) of cells and peptides were incubated in polypropylene tubes for 1hr at room temperature in the dark. To stop the reaction, 30μL of 4% PFA in PBS was added and incubated at room temperature for 10min. To neutralize the fixation, 30μL of neutralization buffer (1.7M TRIS base, 1.25M glycine, pH 9.1) was added for 5min. Intracellular levels of cytochrome C were probed by adding 20μL of staining buffer (1% saponin, 10% BSA, 20% FBS, 0.02% sodium azide in PBS) with 1μL antibody for human cytochrome C. Samples were stained overnight and then transferred into polystyrene tubes for analysis the next day.

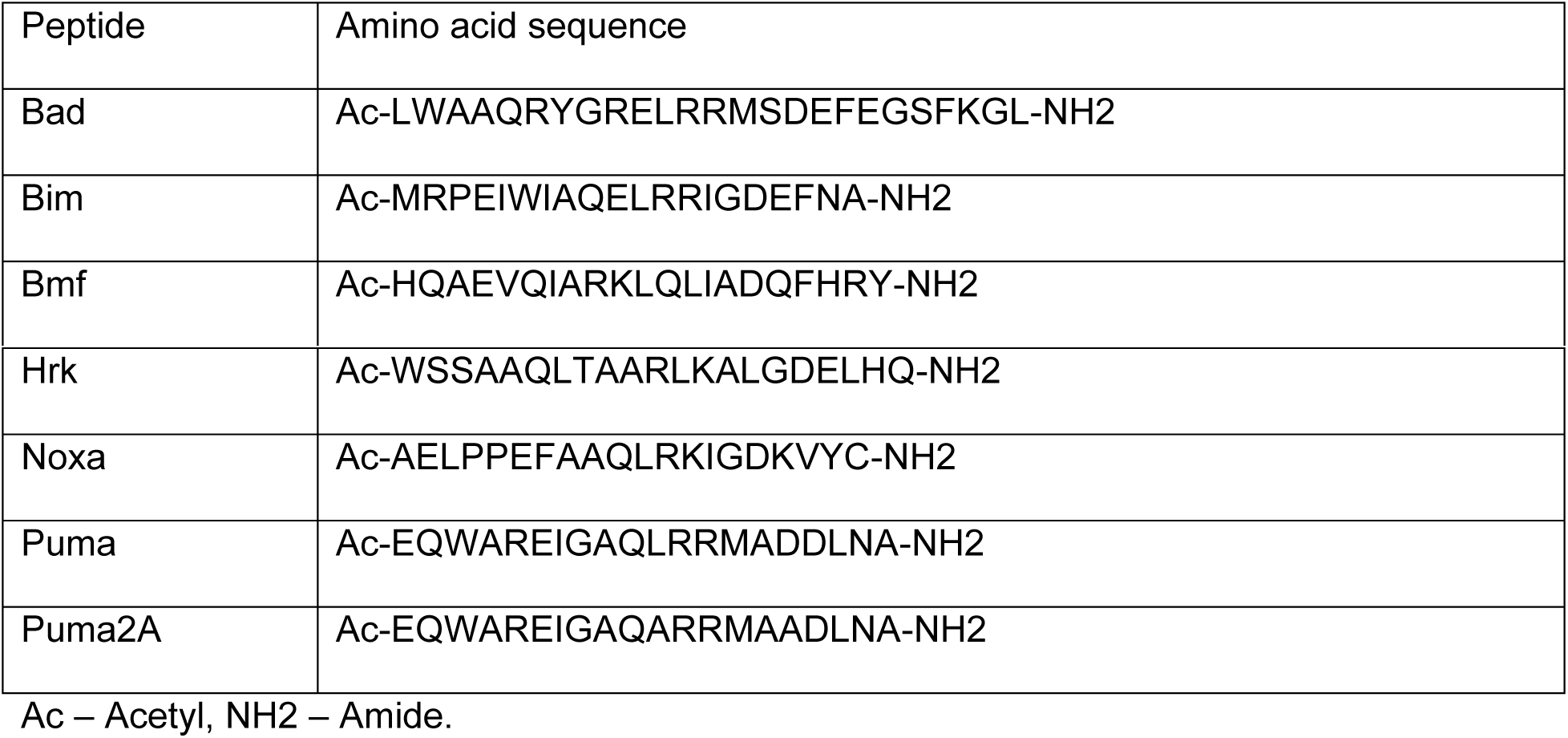

### Flow cytometry

To track proliferation, cells were stained with CellTrace Violet (Invitrogen, C34557). Cells were washed in FACS bufer (5% FBS in PBS), stained with fluorescently-conjugated antibodies for 30min-1hr at 4°C in the dark, and then washed again before being analyzed on a BD FACS Canto II. To normalize acquisition, Spherotech Accucount (ACBP-50-10) beads were included in each tube.

| Antibodies used for flow-cytometry: |  |  |
| --- | --- | --- |
| PE Mouse Anti-Human IgD | BD Pharmingen, 555779 | (no RRID) |
| APC Mouse Anti-Human CD38 | Biolegend, 303510 | RRID:AB_314362 |
| PECy7 Mouse Anti-Human CD19 | eBioscience, 25-0198-42 | RRID:AB_10671548 |
| FITC Mouse Anti-Human cytochrome C | Biolegend, 612304 | RRID:AB_2090159 |

### Dose-response curves and isobolograms

Small molecule inhibitors used in this study were ABT-737 (Selleckchem, S1002), ABT-199 (Adooq Biosciences, A12500), WEHI-539 (APExBIO, A3935), A1210477 (Selleckchem, S7790), and staurosporine (Sigma, S5921-0.5MG). Dose-response curves were generated by treating primary cells for 24 hours at 37°C. Mitogen -activated cells were treated on day 3 post-stimulation for 3 days and assayed on day 6. Cell counts were performed by flow cytometry and normalized to a DMSO (vehicle only) control. Curves were drawn in GraphPad Prism 7 to calculate IC_50_ values and 95% confidence intervals based on a Hill slope factor of 1.

Isobolograms were generated based on dose-response curves with individual drugs (Drug 1, Drug 2) and combinations of drugs (Drug 1 + Drug 2) (Chou, 2010). Individual drug dose-response curves were used to identify drug concentrations at IC_30_, IC_50_, and IC_70_. Similarly, at a constant concentration of Drug 1, the concentrations of Drug 2 were determined at the IC_30_, IC_50_, and IC_70_ points of the combination dose-response curve. To calculate the Combination Index (CI), the following equation was used:

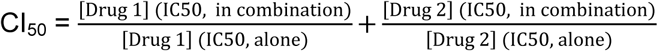

The CI was also calculated for CI_30_ (with IC_30_ values) and for CI_70_ (with IC_70_ values). We classified the interaction between the combination of drugs based on the following CI values:

CI > 1, Antagonistic; CI = 1, Additive; CI < 1, Synergistic.

### Statistical analysis

Student’s 2-way ANOVA and two-tailed t-test were calculated using GraphPad Prism 5.0 software. *p < 0.05 was considered significant. IC50 values were calculated by non-linear regression using GraphPad and 95% confidence intervals were reported.

## Author Contributions

J.D. and M.A.L designed research; J.D. performed research; J.D. and M.A.L. analyzed data; J.D. and M.A.L. wrote the paper.

## Acknowledgements

We thank Lynn Martinek, Nancy Martin, and Mike Cook for extensive help in flow-based cytometry experiments and Karyn McFadden, Alex Price, and Nick Homa for critical reading of the manuscript. Special thanks to Jeremy Ryan from the Letai laboratory for technical help. This work was supported by National Institutes of Health (NIH) Grants R01-CA140337 and R01-DE025994 (to M.A.L.), T32-CA009111 (to J.D.). Additional funding came from the Duke CFAR, an NIH funded program, 5P30-AI064518, and an American Cancer Society grant RSG-13-228-01-MPC (Both to M.A.L.).

---

**Figure 2 Figure Supplement 1.**
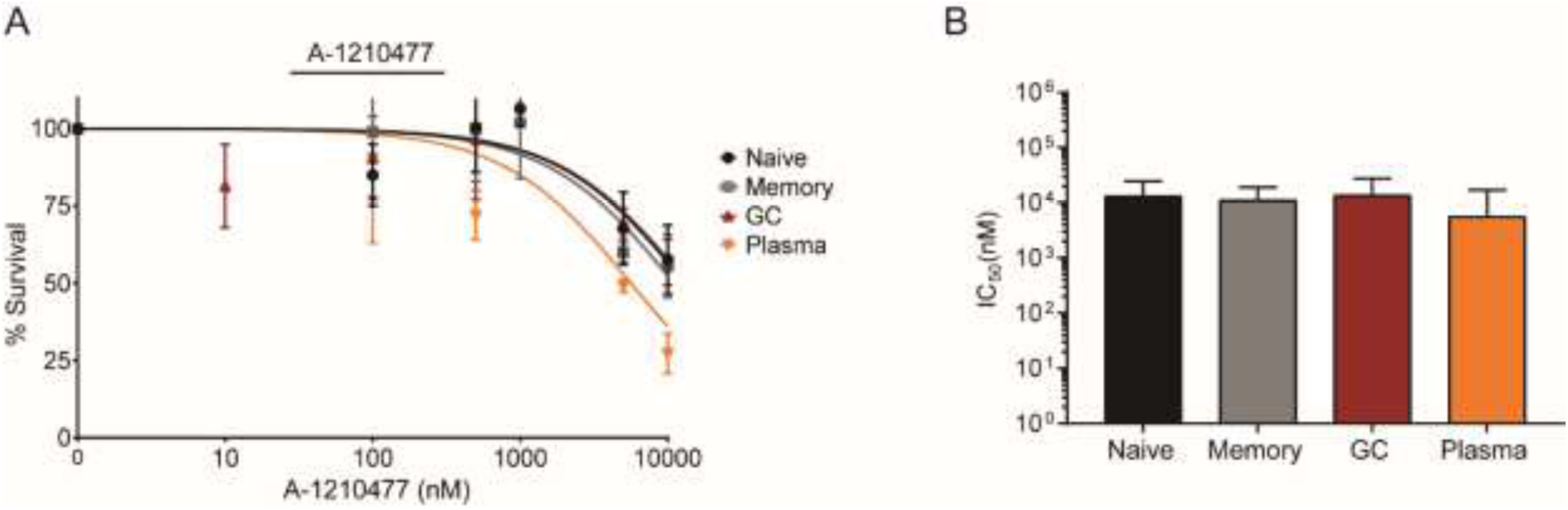
Tonsillar B cell subsets are largely refractory to the MCL-1 inhibitor A-1210477. A) Dose-response curves generated from treating 5 human donors with A-1210477 (MCL-1 specific BH3 mimetic) for 24 hours. B) Average IC_50_ values with 95% confidence intervals for each subset are plotted from (A).

**Figure 4 Figure Supplement 1.**
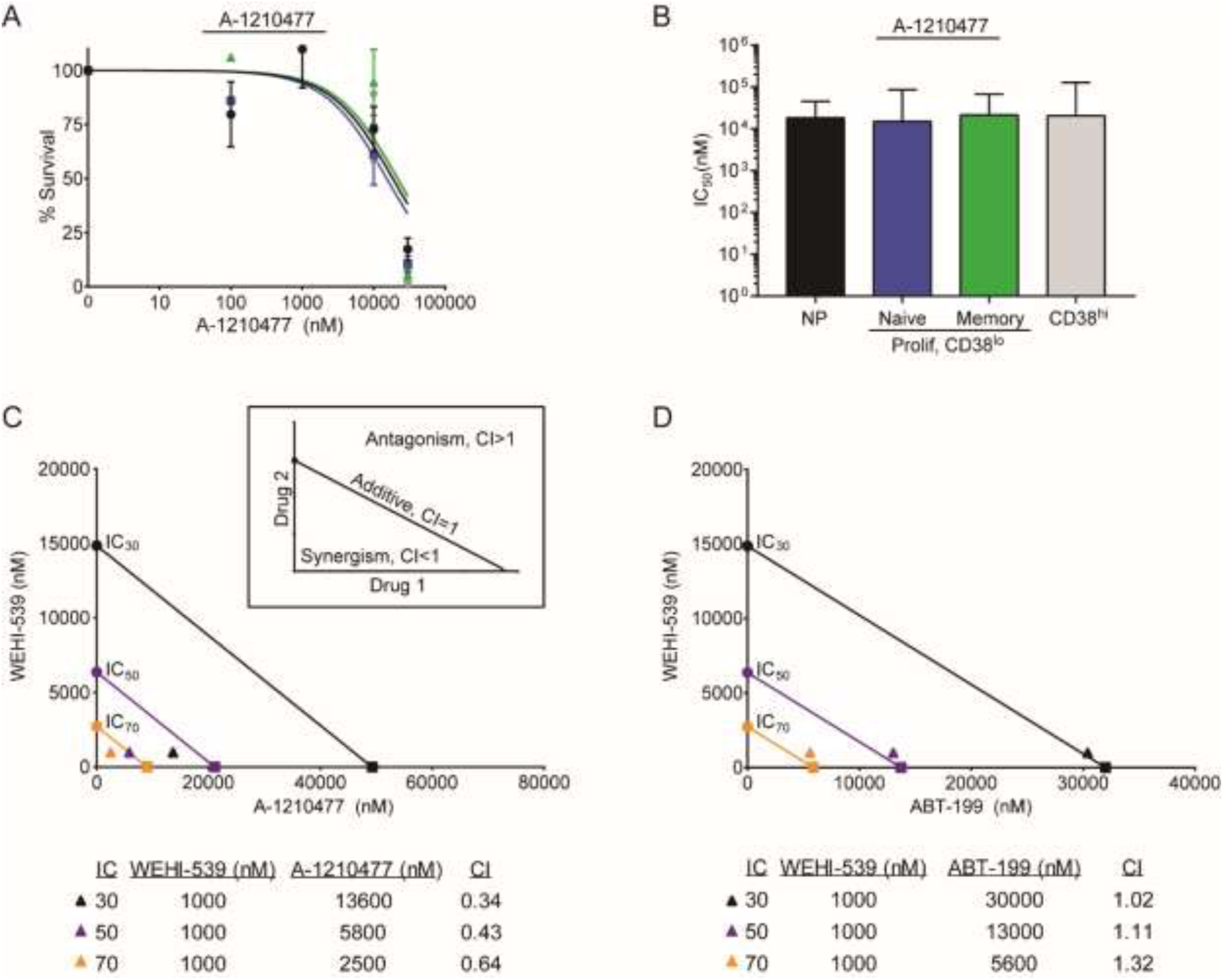
Combined BCL-XL and MCL-1 inhibition is cytotoxic to the CD38^hi^ subset of CpG-stimulated B cells. A) Dose-response curves generated from treating CpG-stimulated PBMCs for 3 days with A1210477 (3 human donors). B) Average IC_50_ values with 95% confidence intervals for each subset are plotted from (A). C) Isobologram generated from treating CpG-stimulated PBMCs for 3 days with WEHI-539 (from Figure 4C), A1210477 (from Figure 4 Figure Supplement 1A), and in combination. IC_30_ (black), IC_50_ (purple), and IC_70_ (orange) values are plotted for WEHI-539 alone (●), A1210477 alone (■), and A1210477 in combination with 1000nM WEHI-539 (▲). Combination index (CI) values are listed below. Inset, a schematic of the combination index (CI) values of synergistic, additive, and antagonistic interactions and where they fall on an isobologram. D) Isobologram generated from treating CpG-stimulated PBMCs for 3 days with WEHI-539 (from Figure 4C), ABT-199 (from Figure 4C) and in combination. IC_30_ (black), IC_50_ (purple), and IC_70_ values are plotted for WEHI-539 alone (●), ABT-199 alone (■), and ABT-199 in combination with 1000nM WEHI-539 (▲). Combination index (CI) values are listed below.

